# Pulsed transistor operation enables miniaturization of electrochemical aptamer–based sensors

**DOI:** 10.1101/2022.06.09.495532

**Authors:** Sophia L. Bidinger, Scott T. Keene, Sanggil Han, Kevin W. Plaxco, George G. Malliaras, Tawfique Hasan

**Affiliations:** Department of Engineering, University of Cambridge, Trumpington Street, Cambridge CB3 0FA, UK; Department of Chemistry and Biochemistry and Center for Bioengineering, University of California Santa Barbara, Santa Barbara, California 93106, United States

**Author notes:** Corresponding author (G.G.M.); (T.H.).

## Abstract

By simultaneously transducing and amplifying, transistors offer advantages over simpler, electrode-based transducers in electrochemical biosensors. However, transistor-based biosensors typically use static (*i*.*e*., DC) operation modes that are poorly suited for sensor architectures relying on the modulation of charge transfer kinetics to signal analyte binding. Thus motivated, here we translate the AC “pulsed potential” approach typically used with electrochemical aptamer-based sensor to an organic electrochemical transistor (OECT). Specifically, by applying a linearly sweeping square-wave potential to an aptamer-functionalized gate electrode, we produce current modulation across the transistor channel two orders of magnitude larger than seen for the equivalent, electrode-based biosensor. Critically, the resulting amplification is scalable, such that there is no signal loss with OECT miniaturization. The pulsed transistor operation demonstrated here could be applied generally to sensors relying on kinetics-based signaling, expanding opportunities for non-invasive and high spatial resolution biosensing.

## Introduction

To address the challenge of signal amplification in electrochemical biosensors, there has been a trend towards transistor-based platforms that both transduce *and* amplify electrochemical signals. Among these, organic electrochemical transistors (OECTs) have emerged as a particularly high-performance candidate due to the large signal gain associated with volumetric ion penetration throughout a conducting polymer channel, commonly composed of the commercially available blend (poly(3,4ethylenedioxythiophene) doped with poly(styrene sulfonate) (PEDOT:PSS) (*1*). By amplifying directly at the sensing location, these systems have been shown to afford better signal-to-noise ratios than simple electrode-based approaches when used, for example, in neural recordings (*2*). OECTs have also been applied to a variety of electrochemical sensing approaches, typically by integrating a biorecognition element at the gate electrode. For example, OECT glucose sensors have been demonstrated in which glucose oxidase is immobilized on a gate electrode such that changes in glucose concentration modulate the source-drain current (*I*_D_) (*3, 4*). Alternatively, gate voltage shifts have also been used to monitor analyte binding in transistor-based biosensors (*5*). All of these applications, however, employ direct current (DC) methods. This works well for systems that supply a continuous signal that is proportional to the analyte concentration (*e.g*., the current produced by enzyme-based sensors, the potentials produced on ion-selective membranes, etc.). However, an important, emerging class of biosensors instead relies on changes in charge transfer rate (*k*_CT_), a signal transduction mechanism that is difficult to monitor effectively using DC methods (*6*).

Among biosensors that employ binding-induced changes in electron transfer kinetics, electrochemical aptamer-based (EAB) sensors are rapidly growing in importance due to their versatility and their exceptional selectivity (*7*–*10*). In these sensors, an aptamer (a DNA or RNA molecule selected in vitro to bind to a specific molecular target) is first reengineered such that target binding causes it to undergo a conformational change (*11*). The aptamer is then modified with a thiol linker for immobilization onto a gold electrode. The opposite end of the aptamer is attached to a redox reporter (typically methylene blue) to generate an electrochemical signal that is independent of any redox activity of the analyte. By altering the distance between the redox reporter and the underlying electrode, the aptamer’s binding-induced conformational change modulates the rate of electron transfer to the electrode (*12*) (Fig. 1), which can be monitored using a variety of transfer-kinetics-sensitive electrochemical methods (*11, 13*). AC voltammetry (ACV), for example, interrogates these sensors using a sinusoidal waveform with a frequency corresponding to the charge transfer rate of interest. The amplitude of the resulting current then depends on the average transfer rate of the redox reporter, thus reporting on the fraction of the aptamers that are target bound (*13*). Because it is even more sensitive to changes in transfer rate, however, square-wave voltammetry (SWV), is now the most commonly used approach to interrogate EAB sensors (*10*). This technique utilizes a square pulse superposed over a linearly sweeping voltage, with the pulse length being tuned to the frequency (transfer rate) of interest and the current being sampled only at the end of each pulse, rendering it particularly sensitive to transfer rates. EAB sensors using SWV interrogation have been shown to support continuous molecular monitoring in situ in the body, including in real-time monitoring of pharmacokinetics and metabolism in awake animals (*14*) and closed-loop feedback control over plasma drug levels (*15*).

**Figure 1.**
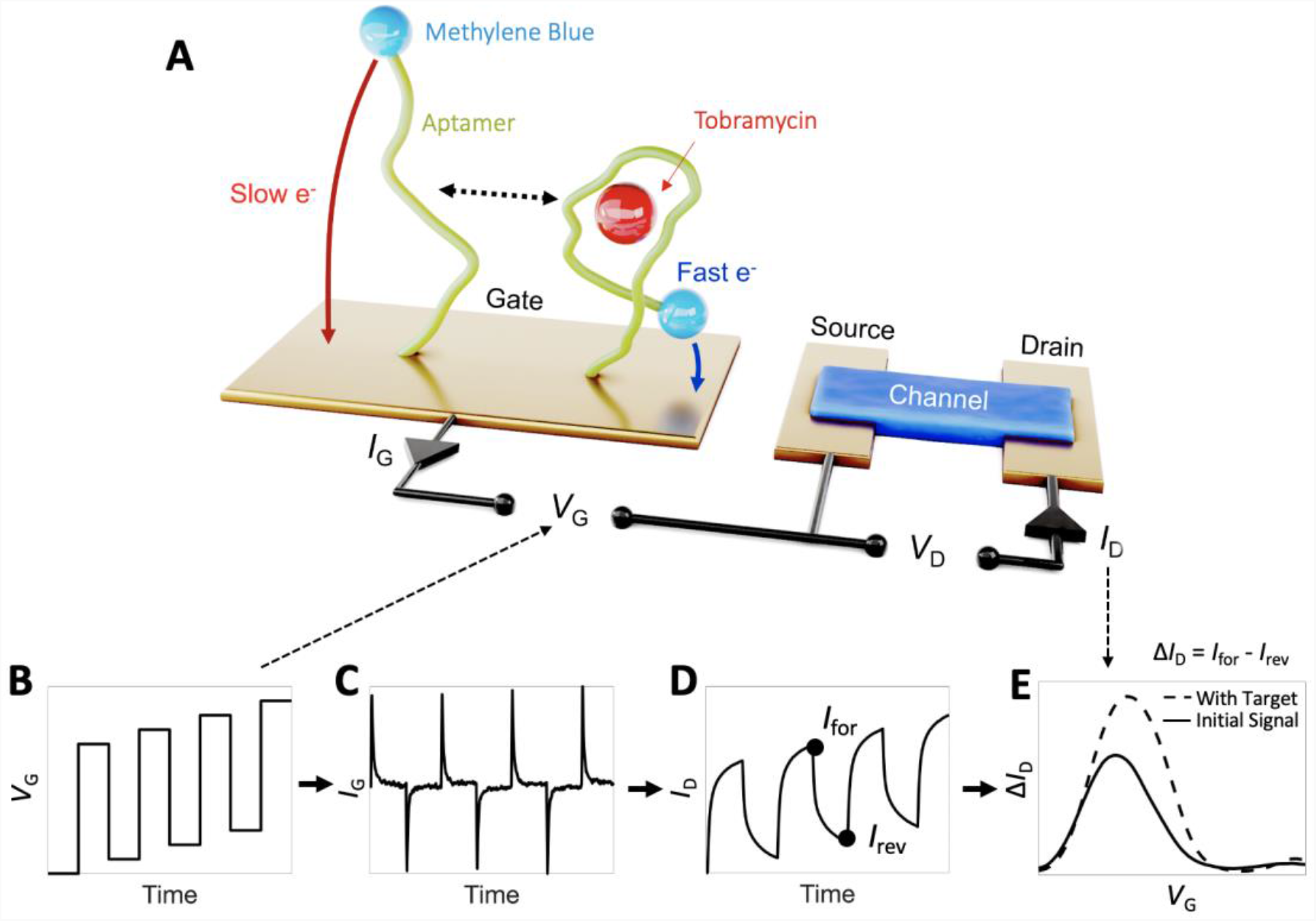
Square-wave gate potential profiles support high-gain aptamer-based OECT sensing. Shown is a (A) schematic of the aptamer-based OECT, which includes methylene-blue-modified aptamers immobilized on the gate electrode. This functionalized planar gold gate and the PEDOT:PSS channel are designed to match capacitance allowing for voltage drop at both the channel and gate sides. **(B)** A pulse square-wave superimposed over a voltage sweep is input as *V*_G_, yielding **(C)** *I*_G_ current decays from the oxidation of the methylene blue. **(D)** The resulting *I*_D_ is proportional to integrated *I*_G_. **(E)** Measuring the difference between each forward and reverse pulse current then yields a distinct methylene blue redox peak. Upon target addition, the charge transfer rate increases, yielding a larger integrated current and leading to a higher Δ*I*_D_ redox peak.

Due to the small currents produced by surface bound aptamers, an increasingly important limitation of traditional (electrode-based) EAB sensors is the difficulty of miniaturizing them below their current, few millimeter length scales (*16*). In response, earlier works have attempted to amplify EAB sensors via OECT-based platforms, but the existing DC transistor sensing methods are not particularly sensitive to changes in electron transfer kinetics, rendering them suboptimal for this application (*17, 18*). Here, however, we adapt square-wave voltage operation at the gate electrode of an OECT to amplify the signals produced by EAB sensors. The resulting amplification is scalable, with two orders of magnitude improvement for planar 0.13 mm^2^ electrodes, suggesting that our approach will enable sensing at high spatial resolution in the mapping of molecular analyte concentrations.

## Results and Discussion

EAB sensors typically utilize a 3-electrode setup, in which a specific voltage (relative to a reference electrode) is applied to the aptamer-functionalized working electrode and the resulting Faradaic (redox-reaction-derived) current flowing to a separate counter electrode is measured. As noted, the transfer kinetics associated with this Faradaic current are typically measured using square wave voltammetry. In this technique, each square voltage pulse drives a current transient decay. By sampling the current at the end of the pulse (the length of which is defined by the square wave frequency), the measured current will be monotonically related to the charge transfer kinetics of the redox reporter. To optimize this strategy, the square-wave frequency is tuned to best distinguish between the transfer kinetics of the aptamer’s bound and unbound states (*11*). The resulting current is correlated to the fraction of aptamers that are target bound, thus reporting on the concentration of the target.

We construct OECT-EAB sensors such that the target-recognizing aptamers are immobilized on the gate electrode (Fig. 1A). Rather than directly measuring the resulting Faradaic current at the gate, however, our approach uses the gate current to modulate the conductivity in the OECT channel. Specifically, the gate current drives ions from the electrolyte into and out of the conducting polymer channel, thereby modulating the charge carrier density (*i.e*., de-doping/doping). Thus, square-wave operation of OECTs is similar to square-wave operation of electrode-based sensors, but with an additional amplification step. In adapting SWV to transistor-based sensors, we apply a square-wave gate voltage (*V*_G_) sweep (Fig. 1B) that yields gate current (*I*_G_) transients (Fig. 1C) caused by electron transfer to or from the methylene blue redox reporter. The OECT amplifies this gate current through the subsequent modulation in source-drain current (*I*_D_, Fig. 1D), which is proportional to the *integrated* gate current:

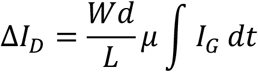

Here *μ* is the hole mobility and *W, d*, and *L* are the channel width, thickness, and length, respectively (*19*).The redox peak is constructed from the differences between *I*_D_ sampled at times corresponding to the end of each *V*_G_ pulse (*I*_for_ – *I*_rev_, Fig. 1D), and the relative peak height reports on the analyte concentration (Fig. 1E).

The gate-to-channel modulation that drives OECT-EAB sensors can be represented by a simple circuit consisting of two capacitors in series, with *C*_G_ representing the gate capacitance and *C*_CH_ the channel capacitance. So as to maintain equilibrium across the circuit, the current flowing at the gate (in response to the redox reactions occurring on it) result in an equivalent charging of the channel (*Q*_G_ = *Q*_Ch_). Such behavior is not usually captured in standard DC OECTs, which are instead designed to maximize voltage drop at the channel side (*C*_G_ >> *C*_CH_) so that the transconductance (*i.e*., voltage-to-current gain, *g*_m_) is maximized. Instead, in our case, we utilize the current gain (*β*) of the OECT to amplify the redox current at the gate.

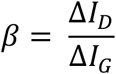

To optimize the signal transduced by the EAB-OECT, we must thus fine tune the gate-to-channel area ratio. Our analysis of various area ratios indicates that a ratio of ∼80 is optimal (*i.e*., at this ratio *C*_G_ = *C*_CH_) (Fig. S1). Lower ratios increase the voltage drop at the gate, increasing the driving force for Faradaic reactions but with lower overall gate currents. In contrast, higher ratios result in too low a voltage drop at the gate to drive the Faradaic reaction as well as increase the background signal from the charging of the electric double layer at the electrode surface, lowering the overall signal-to-noise ratio. Of note, both the device geometry and the drain voltage magnitude affect the redox peak position, which is effectively being referenced against the PEDOT:PSS channel (*5*).

As our testbed to compare our OECT-based EAB sensors with the traditional, electrode-based EAB platform, we focused on sensors employing an aptamer that binds the antibiotic tobramycin. In the absence of this drug, the methylene blue on the aptamer is held relatively far from the electrode surface, yielding a relatively slow charge transfer rate (i.e., small *k*_CT_). Upon binding the drug, in contrast, the aptamer folds into a configuration that brings the redox reporter closer to the electrode, increasing *k*_CT_. Of note, for the electrode-based system, it is possible to select square-wave frequencies at which the redox peak responds to rising target concentrations by either increasing (“signal-on behavior,” which is seen at square wave frequencies that preferentially signal the bound state; see Fig. 2A) or decreasing (“signal-off behavior,” preferentially signaling the unbound state) (*20*). In an OECT-based sensor, however, *I*_D_ is proportional to the *integral* of the current transient, and thus the signal increases with increasing target concentration irrespective of the frequency employed. Specifically, we have empirically found that the maximum signal gain (i.e., maximum relative difference between the bound and unbound states) is reached at 70 Hz (Fig. 2b). This frequency dependence is consistent with a simple simulated model of the aptamer-based OECT system using *k*_CT_ and OECT governing equations obtained from previous literature (Fig. S2) (*14, 19*).

**Figure 2.**
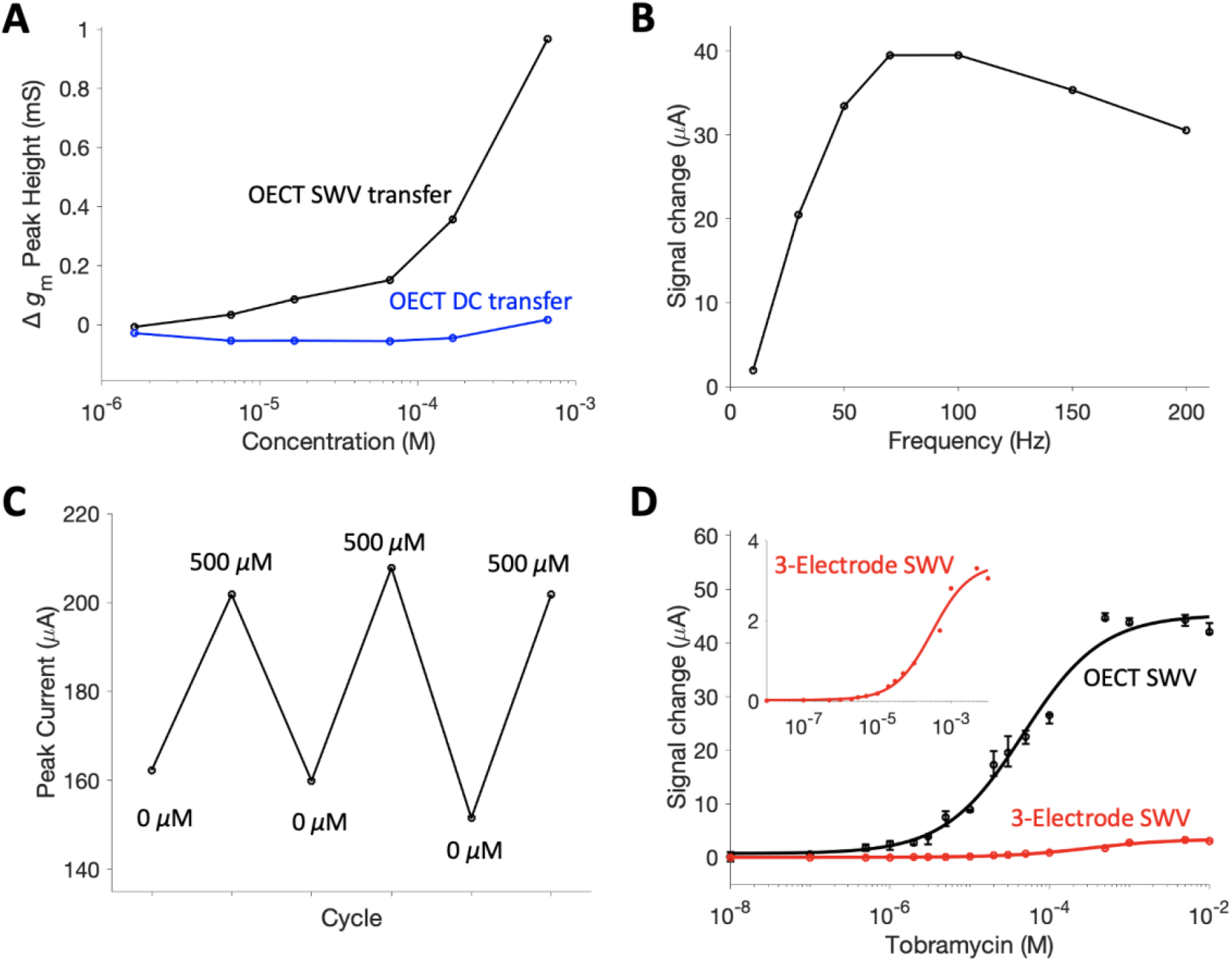
**(A)** EAB OECT sensing behavior showing *g*_m_ peak height growth with concentration using the square-wave OECT operation. The DC transfer curves do not respond to changes in analyte concentration. **(B)** Signal gain upon tobramycin addition versus frequency shows a maximum gain at approximately 70 Hz. **(C)** Peak signal change with spike and rinse cycles indicates good reversibility. **(D)** Binding curves show amplified signal from the OECT current versus 3-electrode SWV using the same gate and working electrode. Error bars correspond to replicate measurements on one device. All data in this figure were measured in 1x phosphate buffered saline (PBS) at ambient temperature. The OECT measurements used *V*_D_ = -300 mV. **(A-C)** correspond to signal gain upon 500 *μ*M tobramycin addition.

The previously reported benefits of EAB sensors are maintained when the approach is adapted to the OECT platform. For example, the excellent reversibility of EAB sensors holds for their OECT implementation (Fig. 2C). Similarly, both the electrode and OECT system exhibit Hill isotherm binding performance (Fig. 2D). The expected Langmuir Isotherm (i.e., saturable) binding has not been demonstrated with DC OECT operation of EAB sensors which may be more sensitive to pH and ionic concentration variation preventing signal saturation. Critically, the concentration-dependent peak height extracted by the AC OECT operation is specific to the electrochemical activity of the aptamer. Because of this aptamer-specific signal, it is suitable for application in complex in vivo environments where selectivity is paramount. To validate this selectivity, we challenged our device in defibrinated horse blood, where its response was quite similar to that seen in simple buffer systems (Fig. 3A).

**Figure 3.**
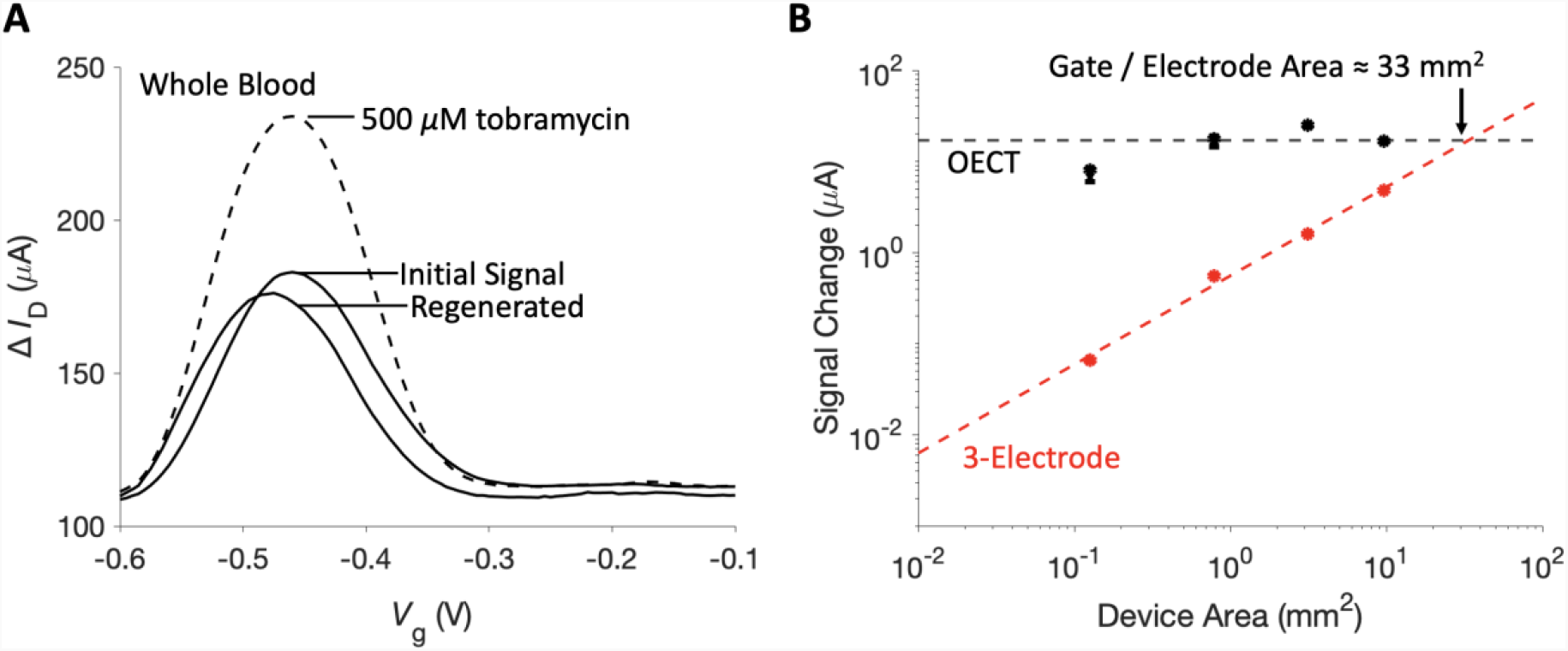
AB-OECT sensors are suitable for in vivo amplification. **(A)** Signal peaks and reversible sensing performance are maintained in whole blood. **(B)** As devices are rescaled for miniaturization, OECT signal is maintained while electrode current decreases linearly with area. Device area corresponds to both the working electrode and gate electrode areas and OECT measurements used *V*_D_ = -100 mV.

A major advantage of applying OECTs in EAB sensors is the ability to maintain an amplified signal even upon substantial miniaturization. The issue is that, in a 3-electrode system, signal shrinks proportionally to reduced electrode area as the electrode accommodates fewer aptamers yielding less absolute current. And while *I*_G_ is similarly reduced in smaller OECTs, due to its lower volume, a smaller channel requires proportionally less current to modulate a change in *I*_D_ (Eqn. 1). That is, as the devices shrink, relative OECT amplification increases and, with this, the absolute signal is maintained. Given this, the OECT-based sensor should yield a higher signal than the traditional 3-electrode system for all electrodes below 33 mm^2^ (Fig. 3B). We believe this highly scalable amplification will allow for a range of sensing applications that were previously infeasible for EAB platforms. For example, miniaturization can enable more sensors per area, which will yield higher resolution spatial measurement.

Beyond the potential future applications of miniaturized EAB sensors, the presented pulsed *V*_G_ technique can be applied generally to a wide range of transistor and biorecognition element pairings. For example, differential pulse voltammetry (DPV) uses a similar waveform to SWV and is used to reduce the effects of capacitive charging and to enable species differentiation by narrowing peaks. While the OECT was selected in this work for its high signal gain and good biocompatibility, the concept is not limited to OECTs; it will operate with the same attributes for any transistor biosensor that signals via a redox event with binding induced *k*_CT_ changes at the gate electrode. By interrogating charge transfer kinetics in this way, pulsed operation will bring the advantages of transistor-derived amplification to a wide range of biosensor approaches.

## Materials and Methods

OECTs were fabricated following a previously reported nanofabrication protocol (Fig. S3) (*21*). First, photolithography with a negative photoresist, AZ nLOF 2035 (Microchemicals GmbH), was used to outline the interconnects and source, drain, and gate electrodes on clean glass substrates. Next, 5 nm of titanium and 100 nm of gold were deposited (Kurt J Lesker PVD-75) after oxygen plasma (Diener Electronic Femto) activation. Gold liftoff was achieved by soaking overnight in NI555 stripper (Microchemicals GmbH). Next, a silane treatment was used for adhesion to the first of two CVD deposited 2 *μ*m parylene-C (PaC) layers (Specialty Coating Systems, Inc.). An anti-adhesive solution was spin-coated before the second PaC layer to enable peel-off. To expose the contact and channel areas, a second photolithography step was completed with a positive photoresist, AZ 10XT (Microchemicals GmbH) and samples were etched with a reactive ion etcher (Oxford 80 Plasmalab plus). The OECT gates and channels were separated by dicing to enable separation for PEDOT:PSS spin-coating. Ethylene glycol (5% v/v) and 4-dodecylbenzenesulfonic acid (DBSA, 0.25% v/v) were sonicated with PEDOT:PSS. Next, 1% (v/v) of (3-glycidyloxypropyl)trimethoxysilane (GOPS) was added and the solution passed through 0.45 *μ*m polytetrafluoroethylene filters. Following another oxygen plasma treatment, two layers of the PEDOT:PSS mixture were spin-coated on the channels for 30 seconds at 3000 RPM. The sacrificial PaC layer was peeled off and the samples were hard-baked to cross link the PEDOT:PSS. Finally, the samples were soaked in DI water overnight and preconditioned to maximize stability (*22*).

The aminoglycoside aptamer (5′−HO−(CH_2_)_6_−S−S−(CH_2_)_6_−GGGACTTGGTTTAGGTAATGA-GTCCC−O−CH_2_−CHCH_2_OH−(CH_2_)_4_−NH−CO−(CH_2_)_2_−methylene blue-3′) was purchased from Sangon Biotech and used as supplied. A 2 *μ*L aliquot of 100 *μ*M aptamer in PBS was thawed and mixed with 4 *μ*L of 10 mM tris(2-carboxyethyl)phosphine (TCEP, Sigma-Aldrich) for 8 hours to reduce the disulfide bond. Next, the aptamer solution was diluted to 500 nM with PBS and placed in a polydimethylsiloxane (PDMS) well over plasma activated gate (working) electrodes for an overnight immobilization. Next, electrodes were rinsed and placed in a 10 mM 6-mercapto-1-hexanol (6-MCH, Sigma-Aldrich) for 5 hours to passivate. Finally, electrodes were rinsed with PBS and connected to channels via a PDMS well.

OECT measurements were collected using a Keysight B1500A semiconductor device analyzer. All OECT measurements used a *V*_G_ sweeping from -600 mV to -100 mV. Electrode measurements were carried out on an Autolab potentiostat (Metrohm) using the gate electrode as working electrode with Ag/AgCl reference and platinum counter electrodes. For square-wave measurements on both the OECT and electrode, a 5 mV step and 20 mV amplitude were used, as adapted from previous work on the aminoglycoside aptamer (*23*). Unless otherwise noted, OECT SWV measurements used a frequency of 70 Hz and electrode SWV measurements used 240 Hz. Sensing performance was measured by adding graduated amounts of stock tobramycin (ThermoFisher).

## Acknowledgments

We gratefully acknowledge figure preparation support from Dr. Ryo Mizuta and Dr. Garam Bae. We acknowledge Dr. Alex Downs for her consultations and guidance regarding EAB sensors.

## Author contributions

The project was initially conceptualized by S.L.B., G.G.M., and K.W.P. Device fabrication and development were carried out by S.L.B., S.H., and S.T.K. Experiments were designed and analyzed by T.H., S.L.B. and S.T.K. S.L.B prepared the manuscript and figures. K.W.P., G.G.M., S.T.K, and T.H. contributed to the writing of the manuscript. T.H., G.G.M., and K.W.P. oversaw the project.

## Data and materials availability

All data are available in the main text or the supplementary materials. Additional data related to this paper may be requested from the authors.

## Supplementary Materials

Supporting information on device geometry, frequency gain fitting, and fabrication protocol available in the supplementary materials.

