## Supplemental figures for "Pulsed transistor operation enables miniaturization of electrochemical aptamer–based sensors"

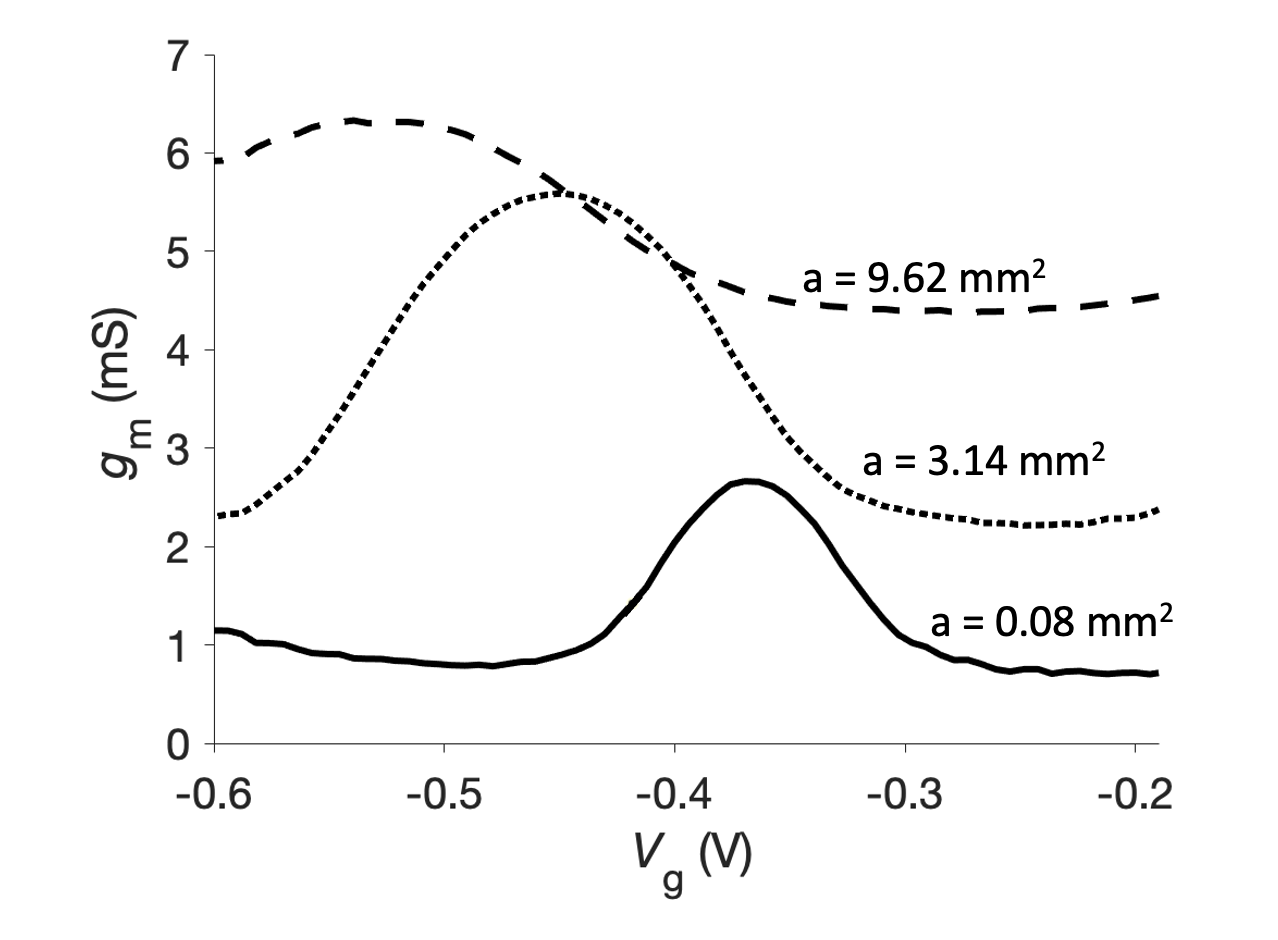

Fig. S1. DC transconductance for various gate electrode areas using the same channel area (W=50 *μ*m, L=400 *μ*m). The methylene blue redox peak shifts to lower voltages as gate area increases resulting in more voltage drop at the channel side.

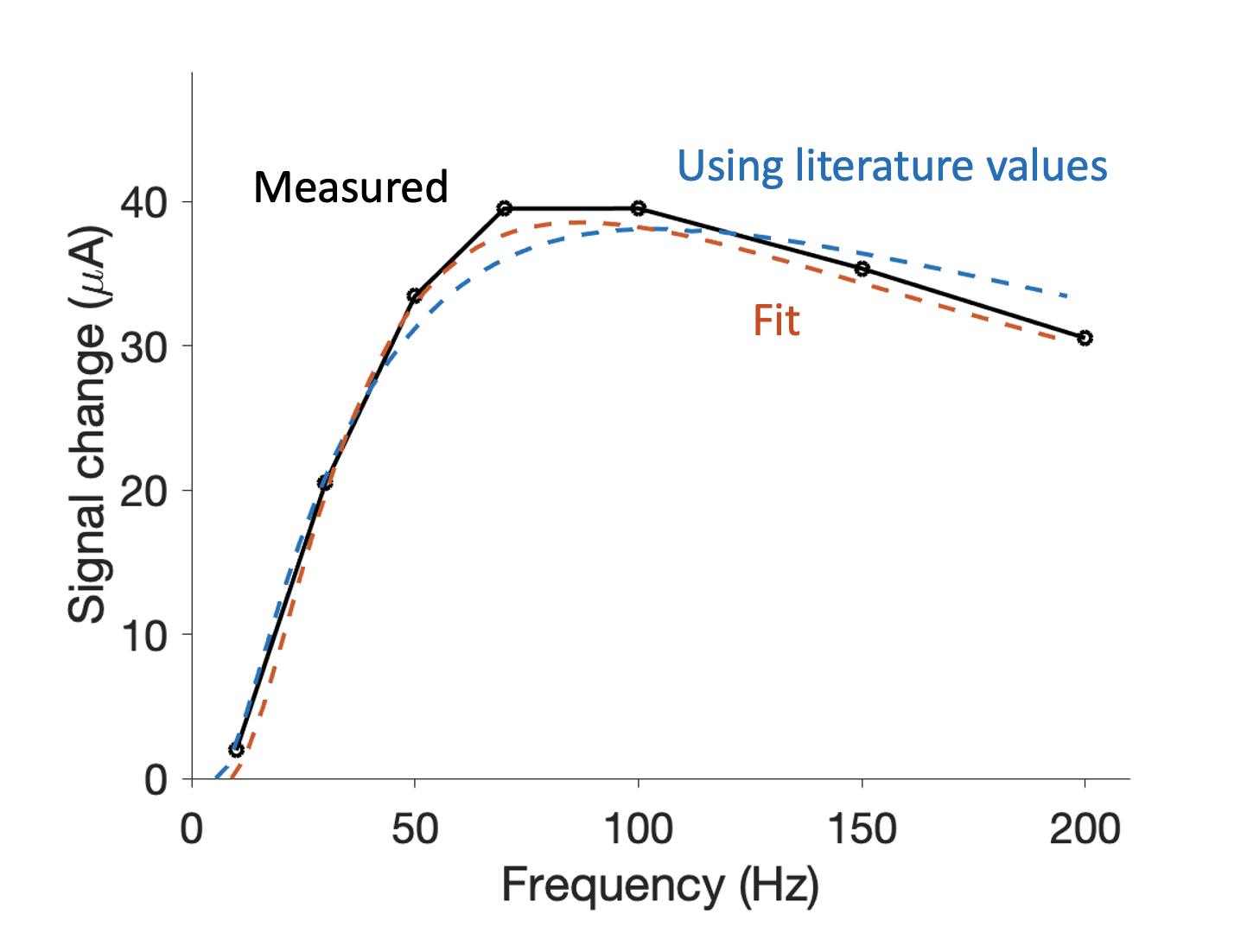

**Fig. S2.** Measured SWV OECT signal gain upon 500 *μ*M tobramycin addition (solid black line) versus expected gain from a basic simulated AB-OECT models (dashed lines). The blue dashed line represents a model using *k*_bound_ and *k*_unbound_ values for the aminoglycoside aptamer from previous literature *(14)* as well as established OECT governing equations *(19)* and the measured aptamer site density. The expected response is calculated from a simplified single square pulse. The orange dashed line corresponds to the same model fit to the measured data, which returns time constants of *k*_unbound_ = 22.3 ms and *k*_bound_ = 6.70 ms.

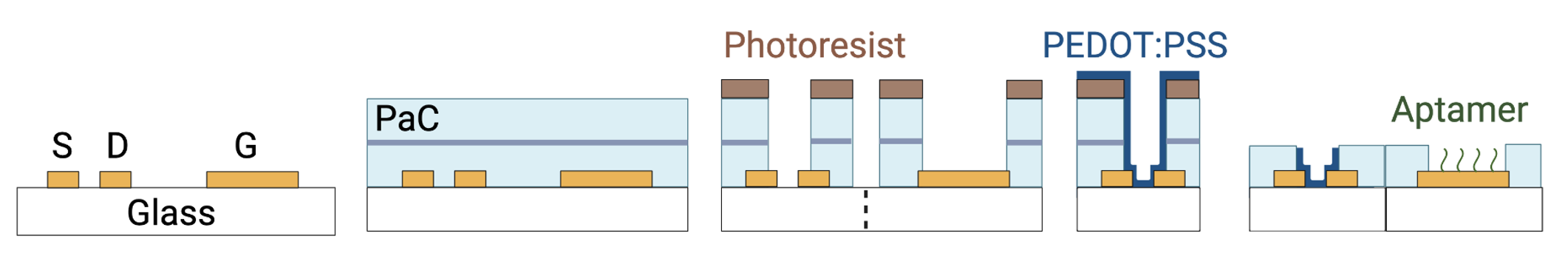

Fig. S3. Aptamer-based OECT fabrication protocol beginning with gold source (S), drain (D), and gate (G) electrode patterning on glass substrates. Next, two parylene-C (PaC) layers are deposited with an anti-adhesive between for peel-off. A second lithography step is used to expose the channel and gate areas then the glass substrates are diced to separate gates and channels. PEDOT:PSS is spin-coated over the channels and the aptamer is immobilized on the gate. The gates and channels are reconnected with a well for testing.
